# Multiscale dynamics and information flow in a data-driven model of the primary motor cortex microcircuit

**DOI:** 10.1101/201707

**Authors:** Salvador Dura-Bernal, Samuel A Neymotin, Benjamin A Suter, Gordon M G Shepherd, William W Lytton

## Abstract

We developed a biophysically detailed multiscale model of mouse primary motor cortex (M1) with over 10,000 neurons and 35 million synapses. We focused on intratelencephalic (IT) and pyramidal-tract (PT) neurons of layer 5 (L5), which were modeled at high multicompartment resolution. Wiring densities were based on prior detailed measures from mouse slice, and depended on cell class and cortical depth at sublaminar resolution. Prominent phase-amplitude-coupled delta and gamma activity emerged from the network. Spectral Granger causality analysis revealed the dynamics of information flow through populations at different frequencies. Stimulation of motor vs sensory long-range inputs to M1 demonstrated distinct intra- and inter-laminar dynamics and PT output. Manipulating PT *I*_h_ altered PT activity, supporting the hypothesis that *I*_h_ neuromodulation is involved in translating motor planning into execution. Our model sheds light on the multiscale dynamics of cell-type-specific M1 circuits and how connectivity relates to dynamics.

---

Understanding cortical function requires studying its components and interactions at different scales: molecular, cellular, circuit, system and behavior. Biophysically detailed modeling provides a tool to integrate, organize and interpret experimental data at multiple scales and translate isolated knowledge into an understanding of brain function. Previous approaches have emphasized structural aspects based on layers and the broad classification of excitatory and inhibitory neurons.^109, 34^ Modern anatomical, physiological and genetic techniques allow an unprecedented level of detail to be brought to the analysis and understanding of cortical microcircuits^79, 1^. In particular, several neuron classes can now be identified based on distinct gene expression, morphology, physiology and connectivity. Excitatory neurons are broadly classified by their axonal projection patterns into intratelencephalic (IT), pyramidal-tract (PT) and corticothalamic (CT) types.^49, 51, 145^ Inhibitory neurons are organized into multiple genetically defined classes, including those expressing Parvalbumin (PV), somatostatin (SOM) and vasoactive intestinal peptide (VIP; also characterized as Htr3A-receptor class).^51, 117, 57^ Recent research has also revealed that connections are cell-type and location specific, often with connectivity differences at different cortical depths within layers.^3, 15, 91^

Primary motor cortex (M1) plays a central role in motor control, but has to date only been modeled to a limited extent.^25, 100, 53^ We and others have extensively studied mouse M1 circuits experimentally, and characterized cell subclasses and many cell-type and sublaminar-specific local and long-range circuit connections.^106, 126, 63^ A major focus of these anatomical and physiological studies has been the distinct cell classes of layer 5 (L5): L5B PT cells – the source of the corticospinal tract, and other pyramidal tract projections, and L5 IT cells which project bilaterally to cortex and striatum. Both of these cell types play a role in motor planning and execution and both have been implicated in motor-related diseases.^125^ Morphology and physiology differs across the two types. L5 IT cells are thin-tufted and show spike frequency adaptation. L5B PT cells are thick-tufted and show little spike frequency adaptation, but strong sag potentials. In terms of their synaptic interconnectivity these types exhibit a strong asymmetry: connections go from IT to PT cells, but not in the opposite direction.^66, 91^ The strength of their local excitatory input connections is also dependent on PT position within layer 5B, with cells in the upper sublayer receiving the strongest input.^3, 55, 143^ These and several other highly specific local and long-range wiring patterns are likely to have profound consequences in terms of understanding cortical dynamics, information processing and function.

We have now developed, and have begun to explore, a multiscale model of mouse M1 incorporating recent experimental data, simulating a cylindric cortical volume with over 10 thousand neurons and 29 million synapses. We attempted, as far as possible, to base parameters on data obtained from a single species, strain and age range, and from our own experimental work. However, these data are necessarily incomplete, and we have therefore combined additional data from multiple other sources. We focused particularly on the role of L5 excitatory neurons, utilizing detailed models of layer 5 IT and PT neurons with full dendritic morphologies of 700+ compartments based on anatomical cell reconstruction and ionic channel distributions optimized to *in vitro* experimental measures. The task of integrating experimental data into the model required us to develop several novel methodological techniques for network simulation design, including: **1.** specifying connections as a function of normalized cortical depth (NCD) – from pia to white matter – instead of by layer designations, with a 100-150 *μm* resolution; **2.** identifying and including specific dendritic distributions associated with particular inputs using features extracted from subcellular Channelrhodopsin-2-Assisted Circuit Mapping (sCRACM) studies ^55, 132^; and **3.** utilizing a high-level declarative modeling tool, NetPyNE, to develop, simulate, optimize, analyze and visualize the model ^37^.

Our model exhibited spontaneous neural activity patterns and oscillations that depended on cell class, layer and sublaminar location, consistent with M1 data. Local field potential (LFP) oscillations in the delta and beta/gamma range emerged, with gamma amplitude modulated by delta phase. Information theoretic measures (spectral Granger causality) showed that information flowed along particular routes at frequencies in the high beta/low gamma band. Different output dynamics were seen when the network was driven by brief activation of particular long-range inputs, or in the setting of different neuromodulatory conditions. The simulations shed new light on how cell-type-specific circuits of M1 are associated with dynamic aspects of activity, including physiological oscillations, neuromodulation and information flow.

## 1 Results

We implemented a biophysically-realistic model of the mouse M1 microcircuit representing a cylindrical volume of 300 *μm* diameter (Figs. 1 and 2). The model included over 10,000 neurons with 35 million synapses. Cell properties, locations, and local and long-range connectivity were largely derived from a coherent set of experimental data obtained using the same techniques in a single lab and single species, with a focus on two deep (L5) populations of particular interest: corticospinal cells and corticostriatal cells. One innovative feature in the network presented here was the inclusion of a Layer 4 for motor cortex, consistent with its recent characterization ^141, 14, 10^. The model was developed using NEURON and NetPyNE ^23, 37^. Over 10,000 simulations were required to progressively construct and improve the model. Simulations required over 4 million high performance computing (HPC) cluster core-hours to arrive at the results shown, primarily during model building. One second of simulation (model) time required approximately 48 core-hours of HPC time. We employed a grid search on underconstrained connectivity parameters – e.g., inhibitory to excitatory weights – to identify simulations that produced realistic activity patterns.

**Figure 1:**
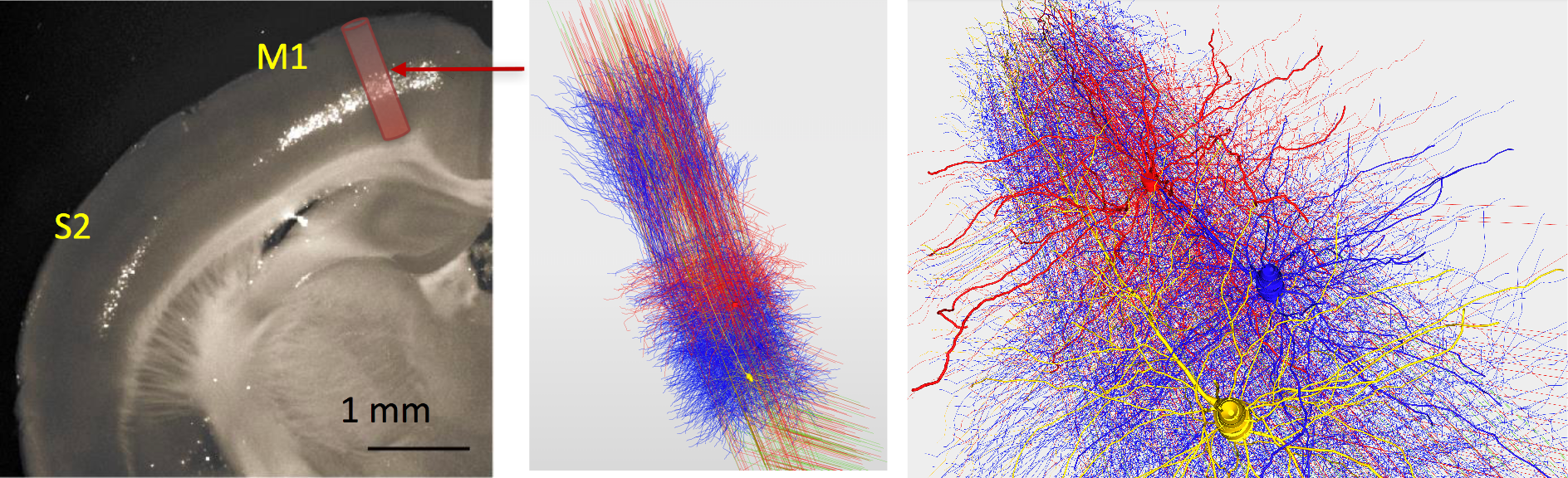
M1 microcircuit model 3D representation (5 % of cells shown). Epifluorescence image of a coronal brain slice of mouse showing M1 and S1 regions, and approximate anatomical location and volume of simulated cylindrical tissue (*left panel*; adapted from ^131^). Representation of M1 network, showing 5% of cells, demonstrates the location and 3D morphologies of model neurons, with one IT (red) and two PT cells (blue and yellow) rendered in detail (*middle and right panels*).

**Figure 2:**
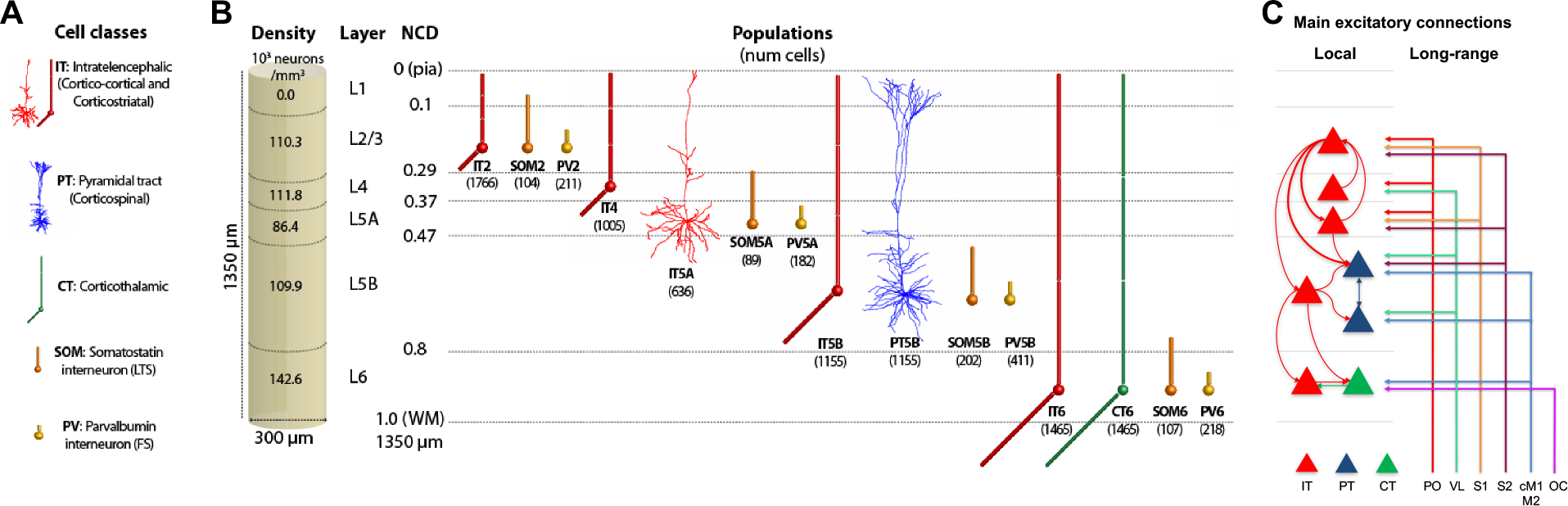
M1 microcircuit model connectivity, dimensions, and neuronal densities, classes and morphologies. **A.** Cell classes modeled. IT5A and PT5B neurons are simulated in full morphological reconstructions. Other excitatory types and inhibitory neurons use simplified models with 2-6 compartments. All models are conductance-based with multiple ionic channels tuned to reproduce the cell’s electrophysiology. **B.** Dimensions of simulated M1 cylindrical volume with overall cell density per layer designation (left), and cell types and populations simulated (right). **C.** Schematic of main local and long-range excitatory connections (thin line: medium; thick line: strong). Note the unidirectional projections from ITs to PTs, with a particularly strong projection arising from L2/3. (IT: intratelencephalic cells–corticostriatal; PT: pyramidal-tract cells–corticospinal; CT corticothalamic cells. PO: posterior nucleus of thalamus; VL: ventrolateral thalamus; S1: primary somatosensory; S2: secondary somatosensory; cM1: contralateral M1; M2: secondary motor; OC: orbital cortex; PV: parvalbumin basket cells, SOM: somatostatin interneurons; number of cells in each population shown in brackets; left shows L1–L6 boundaries with normalized cortical depth–NCD from 0 = pia to 1 = white matter.)

As expected from results in other systems, there was no single “right” model that produced these patterns but rather a family of models (degenerate parameterization) that were within the parameter ranges identified by experiment ^48, 110, 40^. From these, we selected one *base model*, representing a single parameter set, to illustrate in this paper. This base model was tested for robustness by changing randomization settings to provide a *model set*, with analysis of raw and mean data from 25 simulations: 5 random synaptic input seeds × 5 random connectivity seeds (based on connectivity density). The full model set showed qualitatively similar results with low variance in bulk measures (population rates, oscillation frequencies) for changes in randomization settings.

We used the base model and model set to characterize spontaneous firing and LFP patterns in response to activity of long-range inputs. Results are presented both in terms of cell class and cell population. We focused on three excitatory classes: intratelencephalic (IT), pyramidal-tract (PT), corticothalamic (CT); and two inhibitory classes: parvalbumin-staining fast-spiking basket cells (PV), somatostatin-staining, dendrite-targeting, low-threshold spiking cells (SOM). Cell populations are defined by both class and by layer (e.g., IT5A, IT5B indicates class IT in layers 5A, 5B respectively; CT6 is class CT in layer 6). Next, we evaluated model response to short pulses from different regions, and analyzed the dynamical interactions resulting from paired activations. Results are presented in the context of different levels of neuromodulation resulting in changing *I*_h_ conductance in PT cells. We use our results to explain and predict the response of the M1 network to long-range inputs.

### 1.1 Spontaneous activity patterns

In the base model, we characterized *in vivo* spontaneous activity based on expected background drive of ≤5 Hz from all long-range inputs (Fig. 3) ^140, 54^. Excitatory cell time-averaged firing rates ranged up to 25 Hz for excitatory populations, and 60 Hz for inhibitory cell types, consistent with *in vivo* recordings from sensorimotor cortical areas in mouse and rat ^58, 76, 106, 41, 60^. Spiking irregularity (Fig. 3*B*), quantified by mean interspike interval coefficient of variation (ISI CV), was 1.0–2.0 for excitatory neurons and 2.5–3.2 for inhibitory neurons ^27^. Additionally, the distribution of IT and PT cell average firing rates was highly skewed and long tailed, *i.e.,* approximating a normal distribution when plotted in log scale Fig. 3*C*)). The log(*rate*) boxplot (not shown) was symmetric with the median approximately at the center and with symmetric whiskers slightly longer than the subsections of the center box, which is indicative of data coming from an approximately lognormal distribution.^45^ Firing rates observed *in vivo* across hippocampus and neocortex exhibit approximately lognormal distributions ^115, 70, 17, 71, 89^.

**Figure 3:**
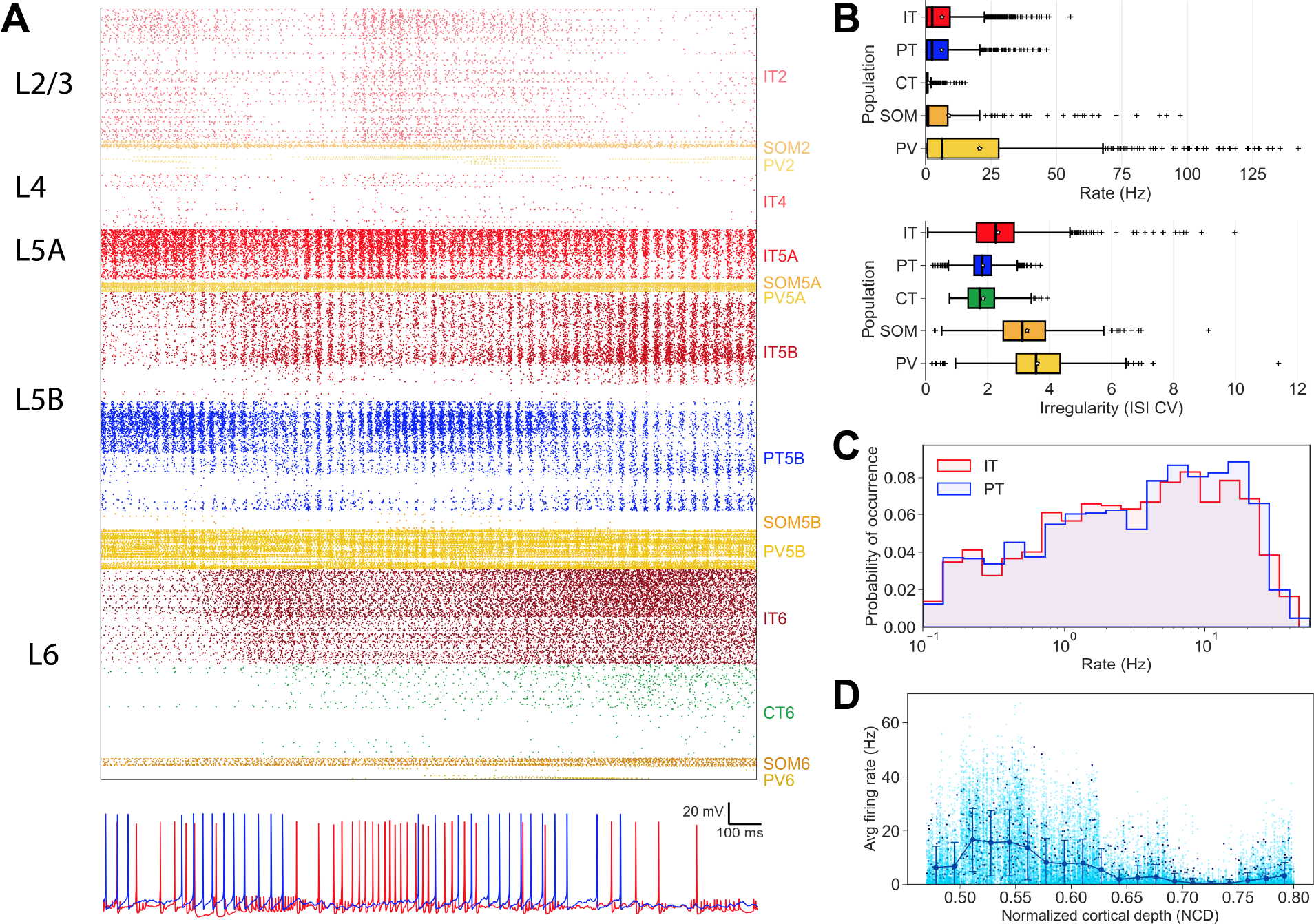
Cell-type and location-specific firing during spontaneous activity. **A.** *Top:* Raster plot of mid-simulation activity (2s of base model simulation shown; cells grouped by population and ordered by cortical depth within each population). *Bottom:* Example IT5 (red) and PT5B (blue) voltage traces. **B.** Firing rates and ISI CV boxplots by cell class in a 50s simulation (IT, PV and SOM cell classes includes cells across all layers). **C.** Distribution of IT and PT firing rates in a 50s simulation. **D.** PT rates by cortical depth shows peak in superficial L5B in full model set of random wiring and inputs (dark blue: base model; cyan: other 24 models).

Activity patterns were not only dependent on cell class and cortical-layer location, but also sublaminar location, demonstrating the importance of identifying connectivity and analyzing activity by normalized cortical depth (NCD) in addition to layer ^51, 3^. For example, PT cell activity was particularly high superficially in L5B, with firing rates decreasing with cortical depth (Fig. 3*D*), consistent with depth-weighted targeting from IT2/3 projections ^3, 139^. This pattern of firing was consistent across network variations with different wiring and input randomization seeds, displaying great variability in firing for individual cells at a given depth (Fig. 3*D* dark blue), but a consistent pattern of rates by depth across all 25 simulations in the model set (Fig. 3*D* cyan). IT5A exhibited similar cortical-depth dependent activity. IT2/3 and IT4 populations showed lower rates than IT5, consistent with weaker projections onto these populations from local M1 ^139, 141^, and from long-range inputs ^83, 132, 141^. In particular, the main source of IT4 input was thalamic, in correspondence with the well-described pattern in sensory cortex ^141^. Within L6, superficial cells of IT6 and CT6 populations were more active than deeper ones. This was due to stronger intralaminar, L5B IT ^139, 142^ (see Fig. 11*A, B*) and long-range inputs, primarily from orbital and contralateral motor cortices (see Fig. 11*C*) ^55^. Weaker local projections onto CT6 compared to IT6 resulted in firing rate differences between CT and IT.

### 1.2 Oscillations and cross-frequency coupling

We simulated recording of local field potentials (LFPs) in the model. LFP signal was calculated by summing the extracellular potential contributed by each segment of each neuron (see Methods for details)^107^. LFP revealed physiologically-realistic oscillations in delta (0.5-4 Hz) and high beta to low gamma (25-40 Hz) ranges across layers and populations (Figs. 4,5). Oscillations occurred in the absence of rhythmic external inputs, emergent from neuronal biophysical properties and circuit connectivity. The large delta oscillation can be readily seen in the raw LFP (Fig. 4*A*). Superimposed fast spikes from local multi-unit activity (MUA) resembled *in vivo* recordings (e.g., ^144^). Filtering the LFP signal from the electrode located in upper L5B revealed phase-amplitude coupling of fast oscillations on delta wave phase (Fig. 4*B*). LFP spectrogram demonstrated that the fast oscillations occurred robustly throughout the time course of simulation (Fig. 4*C*). Beta and gamma frequencies alternated, also coupled to delta phase, with greater gamma at the delta peak, and greater beta at the delta trough (compare Fig. 4*C* with delta wave in 4*B*). We examined LFP phase-amplitude coupling across the model set (N=25) and found a consistently high modulation index ^133^ with average peak frequency for phase at 1 Hz and average peak frequency for amplitude at 40 Hz (Fig. 4*F*).

**Figure 4:**
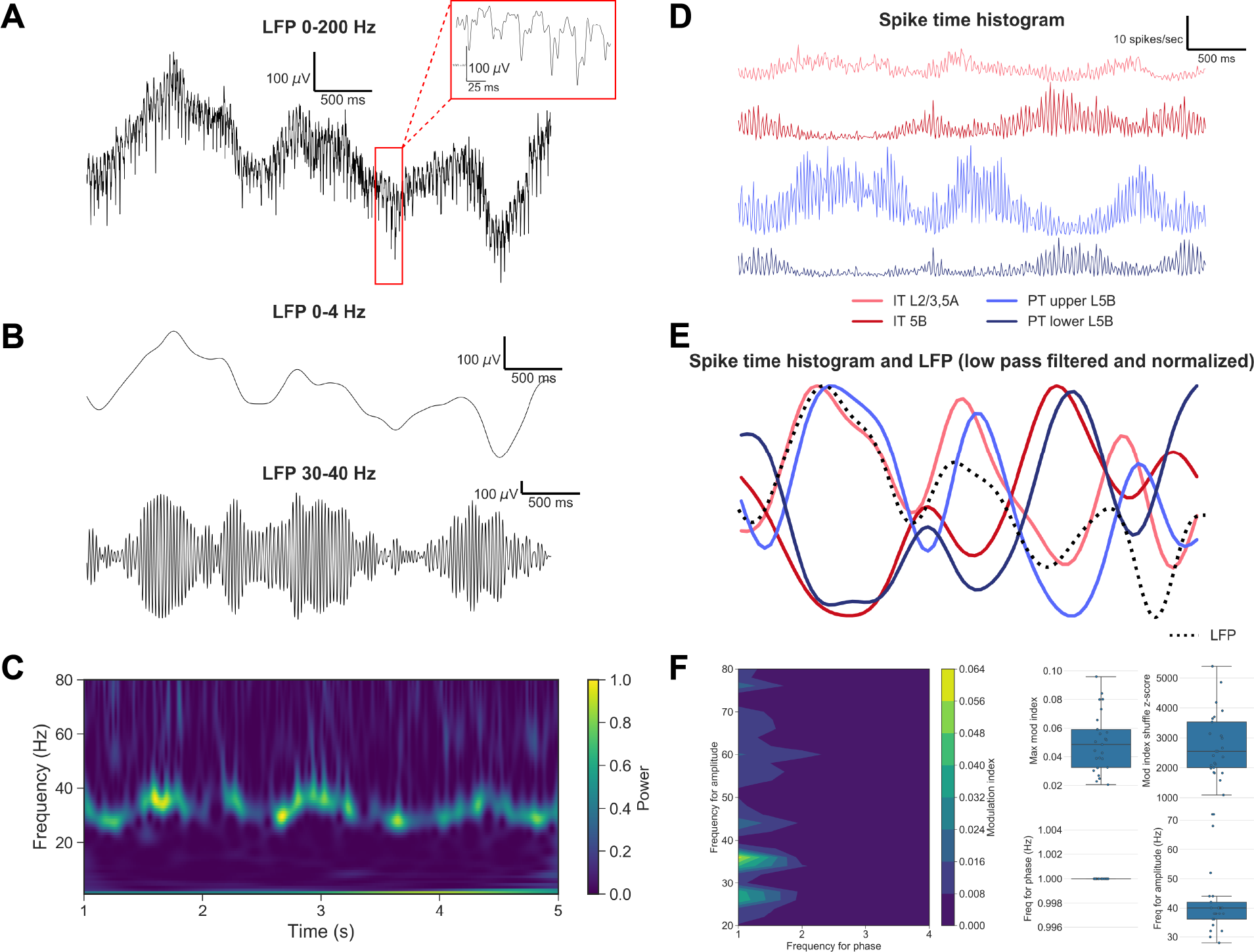
LFP and firing rate oscillations during spontaneous activity. **A.** LFP of base model 4s simulation at L5B (NCD=0.6) at cylinder center. Oscillations are emergent; model has no oscillatory inputs. B. Gamma amplitude modulated by delta phase: 0-4 Hz filtered LFP ~1 Hz delta (top) modulating 30-40 Hz filtered gamma LFP (bottom). **C.** LFP spectrogram: strong high beta/low gamma peak (25-40Hz) modulated by delta phase. **D.** Spike time histogram (firing rate over time) of upper IT (IT2/3 and IT5A), IT5B, PT5B_upper_ and PT5B_lower_. **E.** Upper L5B LFP delta in-phase with IT2/3, IT5A, PT5B_upper_populations and out-of-phase with IT5B, PT5B_lower_populations (low-pass filtered spike time histograms from *D*). **F.** Strong coupling between gamma (36Hz) amplitude and delta (1 Hz) phase from full model set. *Left:* LFP phase-amplitude coupling modulation index for phase at 1-4 Hz (1 Hz step), amplitude 20-80 Hz (2 Hz step) *Right:* Model set statistics showing high modulation indices and significant z-scores (shuffle test) for coupling.

**Figure 5:**
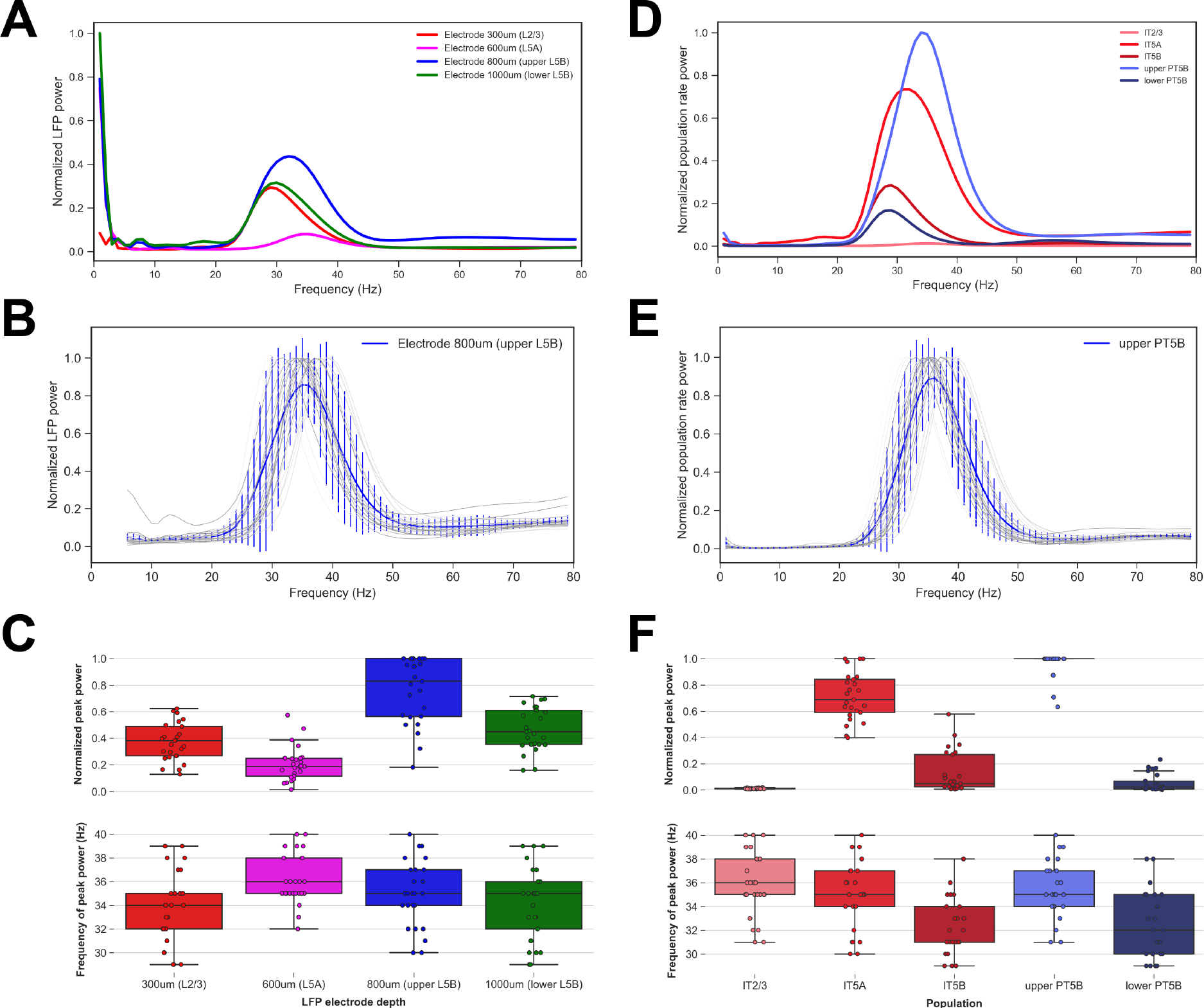
Power spectral density (PSD) of LFP and firing rates during spontaneous activity. **A.** LFP PSDs at different depths for base model simulation: peaks were in high beta / low gamma range (28-35 Hz). **B.** LFP PSDs across full model set (N=25) at upper L5B with mean (blue line) and interquartile ranges (bars). After removing frequencies 1-5 Hz (for clarity), all peak values were between 30-40 Hz. **C.** LFP PSDs statistics of peak power (top) and frequency (bottom) across model set. Average values match trends in panel *A*. **D**. Population PSDs from spike time histograms. Dominant frequencies were comparable to LFPs. **E.** PT5B population PSDs across model set with mean (blue line) and interquartile ranges (bars). All peak values were between 30-40 Hz. **F.** Population PSDs statistics of peak power (top) and frequency (bottom) across model set. Average values match trends in panel *D*.

Both LFP and population rate power spectral densities (PSDs) showed similar beta/gamma oscillatory power coupled to delta phase (Figs. 4*B, E* and 5). Because the LFP is a secondary signal whose phase reverses depending on electrode location, we used the population firing rate delta oscillations to more clearly examine these coupling relationships. IT2/3, IT5A and upper PT5B fired strongly at the delta wave crests and weakly during its troughs, *i.e.,* in phase with delta; L5B IT and lower L5B PT exhibited the opposite pattern, *i.e.,* antiphase with delta (Fig. 4*D, E*). This sheds some light on why the gamma frequency shifts along with delta phase: superficial IT and upper L5B PT cells – active during the crests – oscillate at a higher frequency (~ 35 Hz); IT5B and lower PT5B cells – active during the troughs – oscillate at a lower frequency (~ 29 Hz) (see Fig. 5*D*).

LFP PSD showed evidence that peak frequencies depend on depth (Fig. 5*A*), consistent across different simulation randomizations (Fig. 5*B*). Electrodes in L2/3 (300 *μ*m) and L5A (600 *μ*m) show peak frequencies at 29 and 35 Hz respectively, which could be explained by the influences of the different populations, measured by population firing-rate oscillation frequencies (Fig. 5*D*): ~29 Hz from IT5A, IT5B, lower PT5B versus ~35 Hz from IT2/3 and upper PT5B. The difference between the population-based frequencies and the LFP frequencies can be explained by noting that LFP signals result from synaptic currents so do not directly reflect the population firing at their somatic location ^19, 111^. Instead these synaptic activations drive the population firing. For example, the 29 Hz measured in the superficial electrode reflected postsynaptic currents in large apical and apical oblique dendrites from L5 populations: IT5A, IT5B, lower PT5B. The PT5B population split into two groups with different dominant frequencies, consistent with the upper part of L5B receiving a higher density of intrinsic projections from L2/3 (Methods: Fig. 11*A*). The strong projection resulted not only in higher overall firing rates, but also in distinct modulation of firing (Fig. 5*D*), with higher frequency when the faster-firing upper PT5B cells were dominant, e.g., beginning and middle of Fig. 3*A* raster; and lower frequency when lower PT5B activity increased e.g., towards end of Fig. 3*A* raster. These different frequencies are also partly reflected in the LFPs, where the more clearcut results of population-based activity are smeared by volume conduction ^19, 111^ – for example, L5B LFP includes contributions from inhibitory neurons as well as from both PT and IT excitatory cells. The overall trends in peak amplitude and frequency were conserved across the model set (N=25) for both PSD (Fig. 5*B, C*) and population firing rates (Fig. 5*E, F*), although considerable variability between 28-40Hz was observed for the peak frequencies.

### 1.3 Effect of *I*_h_ modulation on LFP oscillations

We have previously shown that alterations in *I*_h_, modulatable via norepinephrine and other neurohumoral factors ^73^, would alter resonance properties and activation in a model PT5B cell ^98^. Here, we examined whether such alteration in just one cell type would ramify throughout the network to produce widespread dynamical changes. Indeed, reduction in *I*_h_ in PT cells (low-*I*_h_ condition) produced richer dynamics with stronger activity in the gamma band during spontaneous activity (Fig. 6). Higher beta and gamma power was seen in LFPs across all cortical depths, except at 500 and 600 *μm* where oscillatory activity was similarly muted in both low- and high-*I*_h_ conditions. LFP activity across depth also exhibited richer dynamics, with both the amplitude and peak frequency of the beta/gamma oscillations varying over time following the phase of the slower ~1 Hz delta wave, a cross-frequency modulation that was not evident in the high *I*_h_ condition.

**Figure 6:**
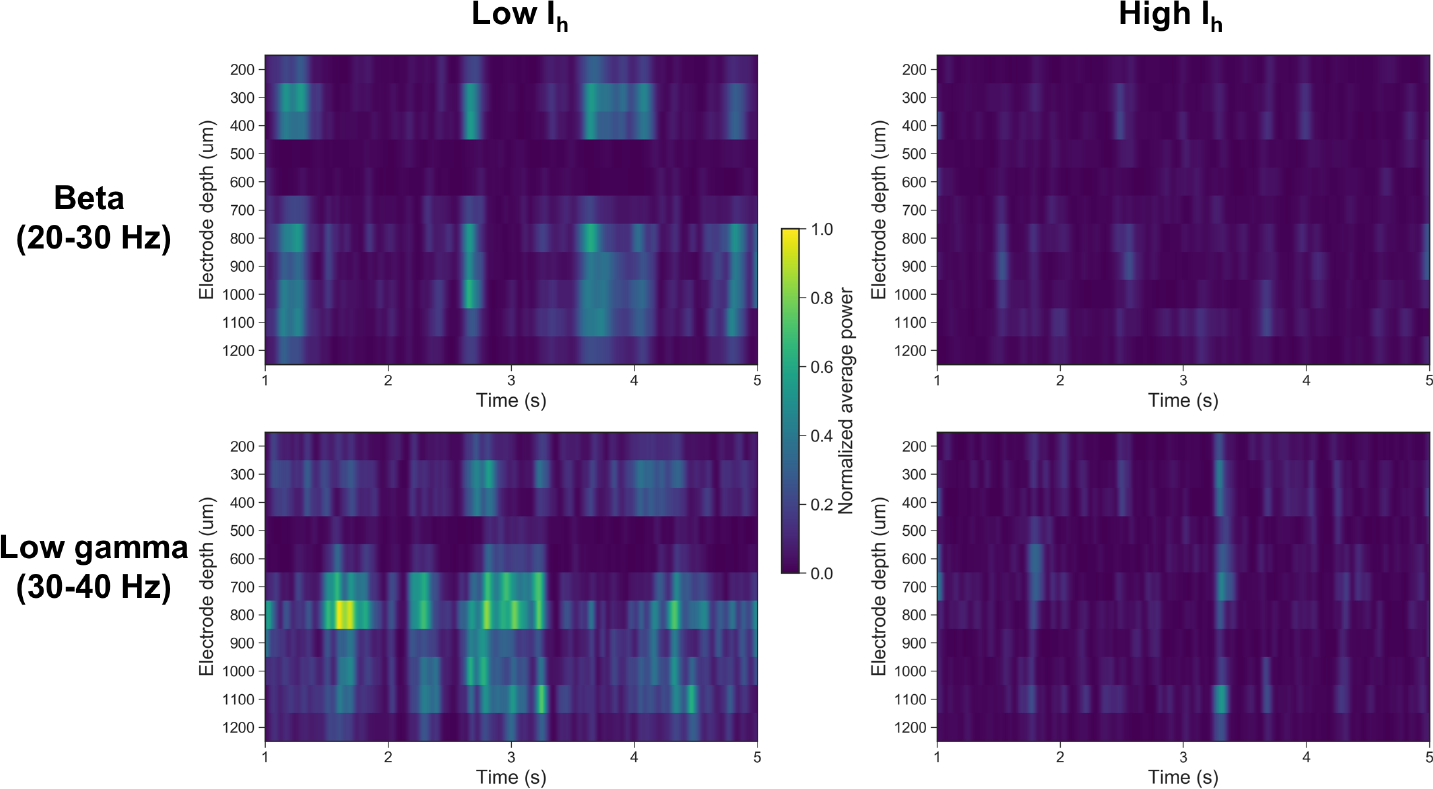
Effect of PT *I*_h_ on network oscillatory dynamics during spontaneous activity. Average LFP power in the beta (20-30 Hz; *top row*) and low gamma (30-40 Hz; *bottom row*) frequency range across electrodes at different depths (y-axis) over 5 seconds. Richer dynamics and stronger beta and gamma are observed with low (*left column*) vs high (*right column*) *I*_h_.

### 1.4 Information flow

Spectral Granger causality (SGC) identified asymmetrical patterns of information flow across the major microcircuit populations (Fig. 7). Information flow in our base model revealed strong SGC at high beta/low gamma frequencies from IT2/3 → IT5A,PT5B, but not in the opposite direction (Fig. 7*A*). Although information flows from IT4 → IT2/3 and IT5A → PT5B were comparatively low, they were considerably higher than flows in the opposite directions. Both IT2/3 and IT5A primarily drove the upper portion of the PT5B population (Fig. 7*B*). The SGC peak for IT2/3 → upper PT5B (31 Hz) was higher than for IT2/3 → IT5A (29 Hz), consistent with the higher dominant oscillatory frequency seen for the upper PT5B vs IT5A populations (Fig. 5*D*).

**Figure 7:**
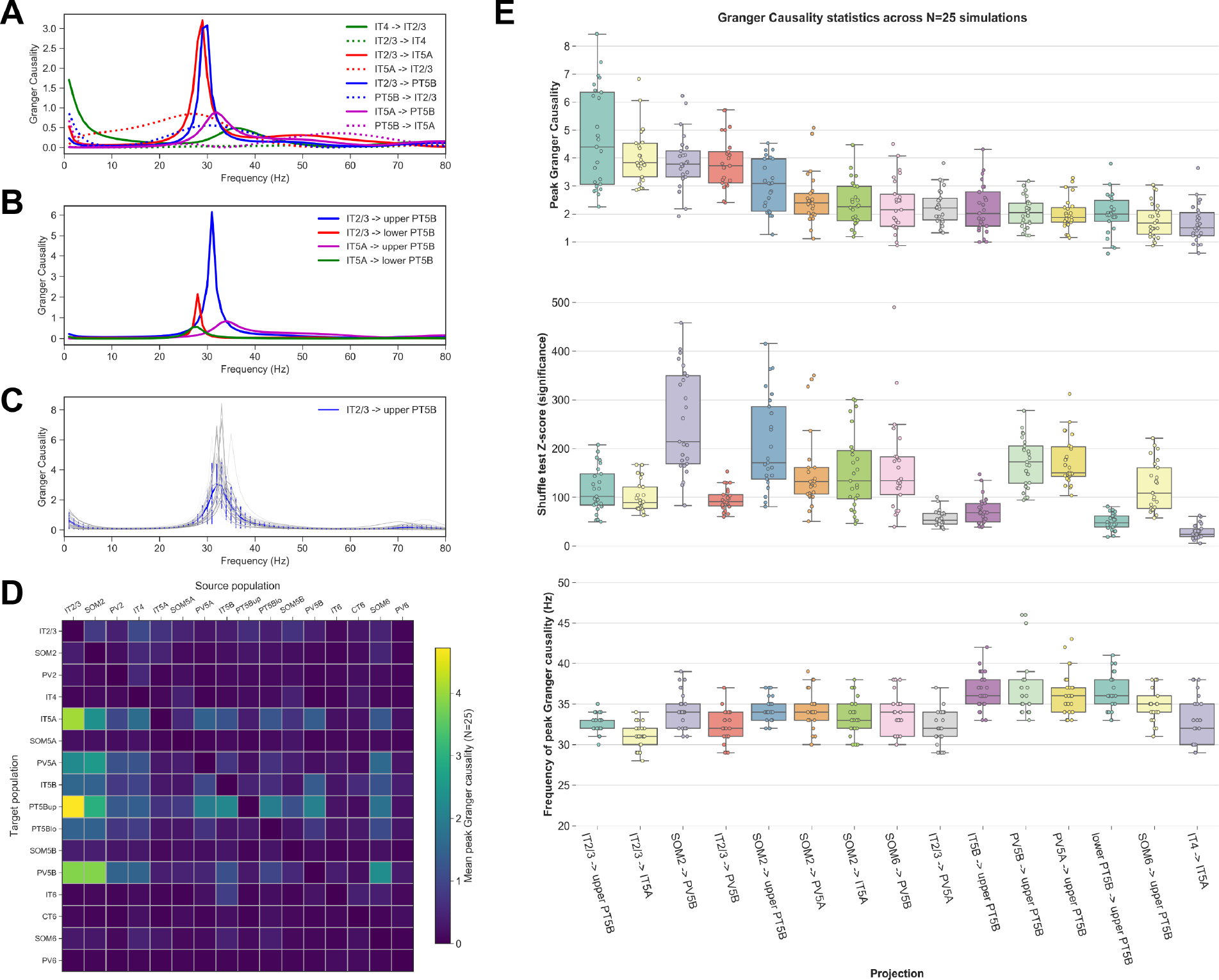
Information flow measure demonstrated by spectral Granger causality (SGC) **A.** Strong SGC in IT2/3→ IT5A; IT2/3→ PT5B; IT4→ IT2/3; IT5A→ PT5B compared to opposite-direction SGC for each case in the base model. **B.** Differential SGC to upper and lower PT5B: strong IT2/3→ PT5B_upper_, compared to IT2/3→ PT5B_lower_. IT5A to both upper and lower PT5B were relatively low (base model). **C.** Comparison of IT2/3→ PT5B SGC across entire model set (gray lines; blue: mean with interquartile ranges). All Granger peak values were between 30 and 35 Hz. **D.** Information flow matrix: peak spectral Granger causality values averaged across model set for all populations pairs. The strongest projections (e.g., IT2/3→ IT5A, IT2/3→ PT5B) match those in the network connectivity matrix (Methods: Fig. 11). **E.** Boxplot statistics of peak Granger causality (top), shuffle test Z-score (middle) and frequency of peak Granger causality (bottom) across model set for the top 15 projections (x-axis).

We analyzed SGC across the model set with their different random inputs and connectivity to investigate robustness of these results (Fig. 7*C*,*D*,*E*). As an example, IT2/3→ PT5B SGC showed a similar profile across the model set, with peaks between 30 Hz and 35 Hz (Fig. 7*C*). We extended the analysis to all pairs of populations (15 × 15 = 225) to create an *information flow matrix* of functional connectivity with mean peak SGC values for each projection (Fig. 7*D*). This analysis revealed strong projections from IT2/3 and SOM2 to IT5A, PV5A, upper PT5B and PV5B, as well as from PV5A, PV5B and IT5B to upper PT5B. We explored the full model set for the 15 matrix projections with the highest peak SGCs (Fig. 7*E*). Although this revealed relatively large variability across amplitudes (Fig. 7*E* top panel), shuffle test z-scores demonstrated peak-SGC significance (mean ≥ 50 except for IT4→ IT5A; Fig. 7*E* middle panel). By contrast with amplitude, peak-SGC frequencies (Fig. 7*E* bottom panel) exhibited relatively low variability. Information flow from IT2/3→ PT5B_upper_ was strongest at a higher mean frequency than IT2/3→ IT5A, consistent with the oscillatory frequencies of the target populations (Fig. 5). All 15 strongest information flow projections targeted L5 cells, and exhibited lower mean peak frequencies when originating from upper layers (L2/3 and L4; 32-34 Hz) than when originating from deeper layers (L5 and L6; 35-36 Hz). This analysis extends the microcircuit description by providing specifics about the cell classes, sublaminar regions and oscillation frequencies. Overall, these results are consistent with the hypothesis of frequency band “labeled lines” differentiating information pathways in the cortical microcircuit ^136^.

Comparing the functional connectivity matrix (Fig. 7*D*) to the anatomical connectivity matrix (Fig. 11) revealed a major overlap but also some differing features. Certain populations had stronger functional influences than would be predicted by anatomy. These included L2/3 IT to L5 PV cells, L2/3 SOM to L5 IT, PT, PV cells, and L5A PV to L5B PT cells. On the other hand, other projections reflected less functional strength than suggested by anatomy: L2/3 PV to L2/3 and L4 IT cells, L5B PV to SOM cells, and L6 PV to IT cells. These differences emphasize the importance of dynamics and multi-synaptic pathways in relating circuit structure to function.

### 1.5 Input pathways and *I*_h_ modulation

Hypothesizing that differences in network dynamics would be reflected in input signal handling, we compared responses to sensory- and motor-related inputs in both high and low PT *I*_h_ conditions (Figs. 8,9). Sensory inputs include similar projections from thalamic posterior nucleus (PO; a higher-order sensorimotor nucleus), and from cortical areas (S1 and S2), all projecting primarily to excitatory cells in superficial layers – IT2/3, IT4 and IT5A (Fig. 8*A*) ^55, 3, 91, 125, 105, 113^. These IT populations in turn project to L5B, where PT cells provide the major M1 output ^3, 132^. Sensory inputs from PO and S2 have also been shown to provide direct, though weaker, projections to PT cells ^55, 132^. We modeled circuit response to a 100 ms sensory input pulse at 10 Hz from PO. This produced an increase in superficial IT and PT5B activity in both low and high *I*_h_ conditions (Fig. 8*B, C*). Increased response in PT5B was mediated indirectly via the strong superficial IT connections, and through weak direct PO projections. PT5B activity was higher for low *I*_h_, both pre- and post-stimulus (Fig. 8*B*). Relative PT5B firing (post-stimulus / pre-stimulus) was similar in both *I*_h_ conditions. This *I*_h_-dependent increase in post-stimulus PT5B response was significant across the full model set (Fig. 8*C*). The increased PT activity may be explained as a consequence of decreased *I*_h_ facilitating synaptic integration of inputs from superficial IT neurons ^123^. CT6, firing rarely, increased activity after stimulation in the low *I*_h_ case. IT5B decreased activity in both high- and low-*I*_h_, potentially as a result of disynaptic inhibition mediated by the increased PT5B response.

**Figure 8:**
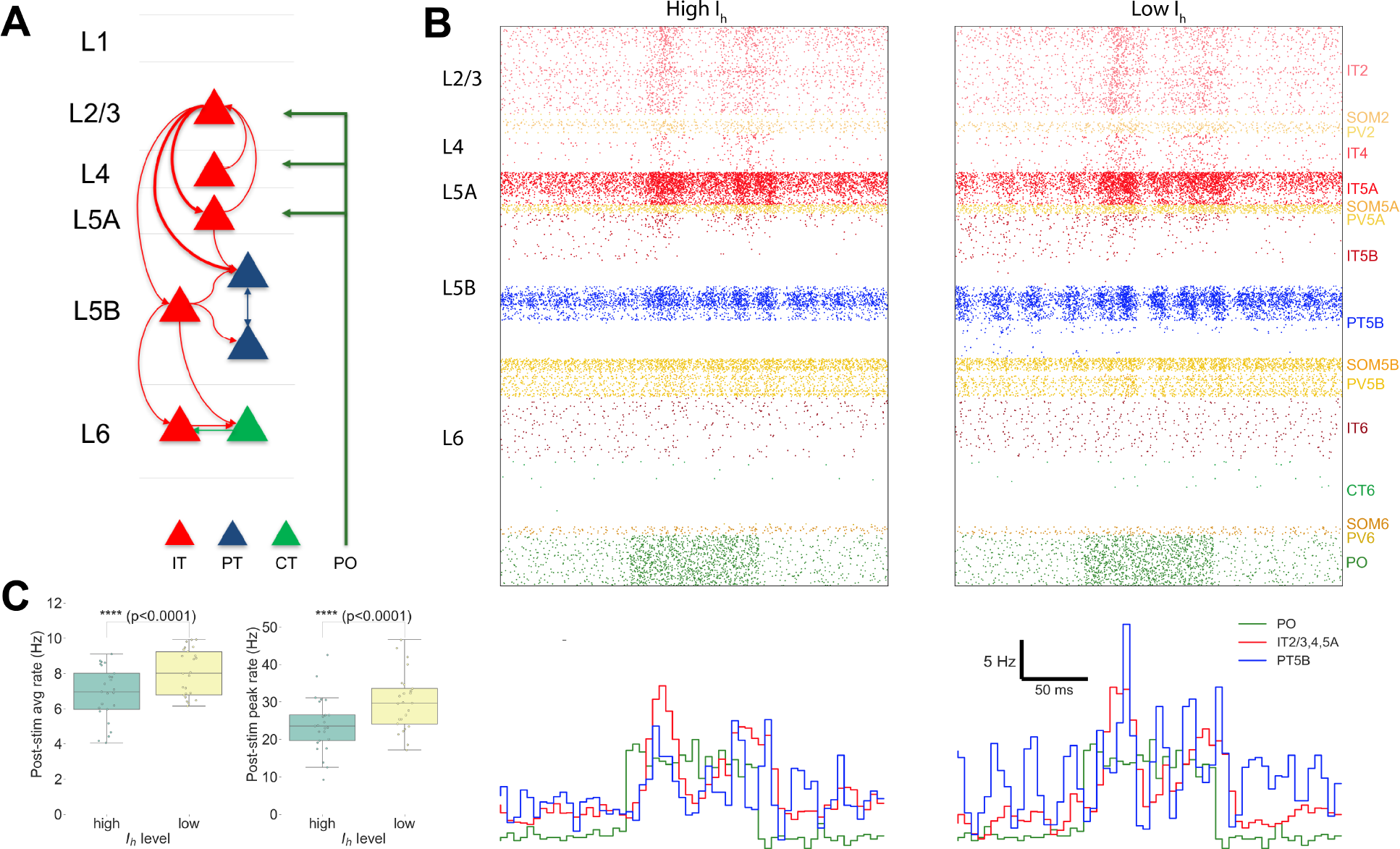
M1 response to input from higher-order sensory thalamus (PO) at high and low PT *I*_h_ levels. **A.** PO input projects primarily to a broad zone of superficial layers, including IT2/3, IT4, IT5A and inhibitory populations. **B.** Raster (top) and spike time histogram (bottom) with 10 Hz PO input (green) for 100 ms with high (left) vs low (right) *I*_h_. Higher activity is seen with low *I*_h_. **C.** PT5B showed higher post-stimulus average and peak firing rates for low PT5B *I*_h_ across model set.

**Figure 9:**
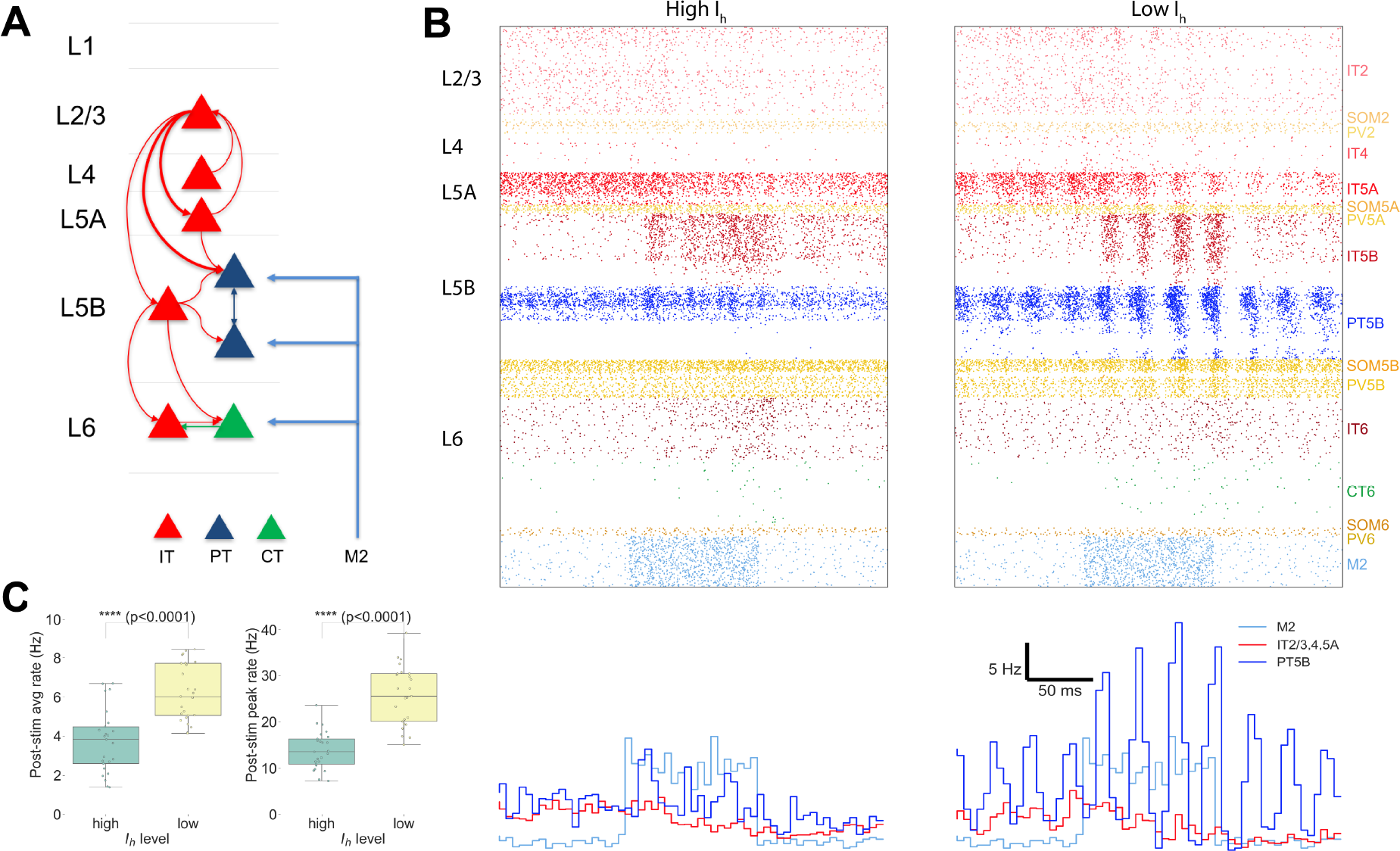
M1 response to input from motor-related inputs (M2) for high vs. low PT *I*_h_ levels. **A.** M2 input to deep layers directly activates PT5B and CT cells, and other cell types in the indicated layers, including inhibitory cells. **B.** Raster (top) and spike time histogram (bottom) of response to M2 input (blue) of 10 Hz for 100 ms: high (left) vs low (right) PT5B *I*_h_. M2 input resulted in higher superficial IT and PT activity, with PT5B exhibiting a stronger response for the lower *I*_h_ case. **C.** PT5B showed higher post-stimulus average and peak firing rates for low PT5B *I*_h_ level in full model set.)

By contrast with the superficial layer projections of sensory-related inputs, motor-related inputs (ventrolateral thalamus, VL; and motor cortical areas, cM1 and M2) project strongly to deep layers and therefore directly excite the output (PT5B) cells (Fig. 9) ^3, 132, 55^. Motor inputs elicited a PT5B response that was similar to that elicited by sensory inputs in the *I*_h_-high condition. However, the response in the *I*_h_-low condition was stronger and longer-lasting than seen with the sensory-related input. Specifically, the stimulation-related 35 Hz PT5B oscillation in the low *I*_h_ setting was more robust in the motor condition, outlasted the duration of the stimulus, and showed spread to IT5B and IT5A despite less overall IT5A response (compare Fig. 8*B* with 9*B*). The increase in both average and peak post-stimulus firing of PT5B cells for low vs high *I*_h_ was also statistically significant in this case (Fig. 9*C*). Weaker PT5B response to sensory inputs compared to motor inputs may be partly due to the initial pulse activation reducing as it propagated from superficial layers to the deeper PT5B cells.

## 2 Discussion

Our ambition was to develop a detailed multiscale computational model of the mouse M1 microcircuit. We necessarily fell short due to lack of data on a number of key molecular, cellular, network and external connectivity aspects, as will be described in *Limitations* below. Despite this, we believe that our model constitutes the most biophysically detailed model of mouse M1 microcircuit currently available. In terms of analysis, our focus has been on the internal dynamics driving cortical output pathways from L5. We therefore have put the greatest effort into the modeling of the two types of L5 projection neurons: the intratelencephalic (IT) and pyramidal-tract (PT) neurons (Fig. 2). Overall, our model incorporated quantitative experimental data from 30 studies, with 11 of these coming from our experimental laboratory, and 19 from other labs. Model development also benefited greatly from extended discussions between the computational and experimental authors.

We used the model to explore a set of questions regarding emergent dynamics, as well as neuromodulatory and dynamical control of input-output relations. The firing rate range generated by our model for each cell type (Fig. 3) compared favorably to experimental results ^115, 70^, and to other recent models ^109, 85, 6, 121^. Fast-on-slow oscillations emerged across multiple random wiring seeds and background input seeds (Fig. 4), a characteristic oscillatory pattern found in biology ^21, 8^, and in neural models ^114, 96^. LFP frequencies varied somewhat across cortical depth. A range of frequencies could also be assessed across individual cell populations (Figures 5*D* and 4*D, E*). Similar frequencies emerged as peaks in measures of Granger causality from one population to another (Fig. 7).

Consistent with the anatomy of projections into M1,^55, 132^ we investigated long-range inputs targeting superficial and deep layers. Response dynamics differed across these two input types, and were also altered substantially by neuromodulatory state (Fig. 8,9). This revealed how these two different functional pathways could generate distinct corticospinal output. Sensory-related inputs from PO, S1 and S2 regions, projecting primarily to superficial IT neurons, produced secondary activation, as well as weak direct activation, in layer 5 corticospinal cells. The deep motor-related inputs from VL, cM1 or M2 regions bypassed the upper M1 layers, projecting directly to the output layer and producing a more immediate and more robust response in the corticospinal cells. *I*_h_ downregulation increased corticospinal response to both types of inputs.

### 2.1 Limitations

This model was constructed over a period of three years. Over the course of that time we updated the model periodically as data came in, but of course any neurobiological model is always in need of continuous updating and improvement as new measurements become available. The situation is comparable to weather models which are gradually improved over months and years even as they are daily updated with current initial conditions to provide forecasts.

Of some concern is the relative lack of data on dendritic ion channel density, which will affect the influence of distal synaptic inputs on L5 neurons. Cell models are precisely tuned to reproduce experimental somatic responses, but limited data is available to characterize dendritic physiology. We have also neglected within-class cellular diversity in our model – all the model neurons of the same cell class have identical morphologies and identical channel parameters. This contrasts with other models which vary both channel conductances and morphologies, the latter by slightly jittering angles and lengths.^84^

Due to the nature of our LSPS and sCRACM data, our model used connection density based on postsynaptic cell type and presynaptic locations. Our model’s normalized cortical-depth-dependent connectivity provided greater resolution than traditional layer-based wiring, but still contained boundaries where connection density changed and did not provide cell level point-to-point resolution. Other recent models have used a sophisticated version of Peters’ principle (identifying overlap between axonal and dendritic trees) to provide cell-to-cell resolution for selected cells, which must then still be replicated and generalized across multiple instances to build a large network ^112, 84^.

We are limited not only by lack of precise data for parameter determination, but also by computational constraints. Often, network simulations use point neurons in order to avoid the computational load of multicompartment neurons, but at the expense of accuracy. Here, we compromised by using relatively small multicompartment models for most populations, with the exception of the neurons of L5. In terms of norepinephrine influence, we focused here on one effect on once cell type, neglecting its wide-ranging effects ^103^, as well as the influence of second messenger cascades ^95^. Even with these compromises, optimizing and exploring our large network model required millions of HPC core-hours.

### 2.2 Emergence of physiological oscillations and phase-amplitude coupling

Our model of M1 neocortex exhibits spontaneous physiological oscillations and cross-frequency coupling without rhythmogenic synaptic input. Strong oscillations were observed in the delta and beta/gamma ranges with specific frequency-dependence on cell class and cortical depth (Figs. 4 and 5). Strong LFP beta and gamma oscillations are characteristic of motor cortex activity in both rodents ^24, 136^ and primates ^116, 102^, and have been found to enhance signal transmission in mouse neocortex ^127^. Both beta and gamma oscillations may play a role in information coding during preparation and execution of movements ^2, 136^. More generally, these physiological oscillations are considered to be fundamental to the relation of brain structure and function ^20^. Phase-amplitude coupling of delta/beta and theta/gamma oscillations has also been reported in motor and somatosensory cortices of rodents ^94, 59, 144, 4^ and primates ^118, 32, 90^. Cross-frequency coupling may help integrate information across temporal scales and across networks ^21^.

As the primary output, PT cells receive and integrate many local and long-range inputs. Their only local connections to other L5 excitatory neurons are to other PT cells ^66^. However, by targeting inhibitory cells in L5,^5^ they are able to reach across layers to influence other excitatory populations, either reducing activity or entraining activity ^93^. These disynaptic E→I→E pathways likely play a role in coupling oscillations within and across layers, and in setting frequency bands.

### 2.3 Information flow

Spectral Granger causality (SGC) measures the ability of a time series X to predict the future values of time series Y, beyond what is possible just with past values of Y. In our context, SGC can be best described as a measure of information flow.^130, 26^ Granger causality estimates, like other information theoretic measures, may be severely biased or show high variance, leading to spurious and potentially misleading results.^130^ We partly addressed these concerns using shuffle-test z-scores, which showed high significance for SGC measures compared to the data without the original time stamps. We were also able to demonstrate fairly low variability across our model set of 25 randomized simulations (Fig. 7*C*,*E*).

Here we provide, for the first time, a full matrix of Granger causality across all projections in a biophysically detailed circuit model (Fig. 7*D*). SGC analysis provided a tool to identify sub-networks of importance, and to reconstruct functional networks that can be contrasted with the underlying anatomical network. The majority of functionally strong projections were also present in the anatomical connectivity matrix, suggesting how dynamics could be used to infer network wiring. For example, the projections from L2/3→L5A and L2/3→L5B could be independently identified. Interestingly, some multi-step pathways which could be identified anatomically showed evidence of not being frequency-coherent dynamically. In this way, dynamical analysis can provide clues as to how different projections could be concatenated, kept separate, or embedded, in the context of the labeling-by-frequency hypothesis ^136, **?**^.

Although functional connectivity largely followed structural excitatory connectivity, there were exceptions that could be partly explained by the strong influences of multiple inhibitory interneuron populations. SOM2, SOM6, PV5A, PV5B influenced both excitatory neurons and other inhibitory neurons. These effects could be the source of interlaminar oscillatory coupling and synchronization through both multi-synaptic inhibition and disinhibitory mechanisms. For example, considering just the strongest projections, PT5B are inhibited by PV5A, PV5B and SOM2, but SOM2 also disinhibits PT5B via the SOM2 → PV5B → PT5B chain. At the same time, the strong IT2/3 → PT5B pathway has a parallel disynaptic inhibitory chain IT2/3 → PV5B → PT5B. Statistical analysis of the strongest projections further revealed different information flow peak frequencies for superficial vs deep layers of origin (Fig. 7*E*). Further analysis of these multiple interacting frequency-based interactions will require development of new statistical and machine learning techniques.

### 2.4 Effect of HCN current (*I*_h_) in PT cells

The concentration of HCN channels has been shown to be significantly higher in PT cells than in IT cells.^123, 52^ Downregulation of *I*_h_, effected via norepinephrine and other neuromodulatory factors, has been shown to increase PT activity as a consequence of enhanced temporal and spatial synaptic integration of EPSPs.^123, 73^ Paradoxically, *I*_h_ downregulation has also been reported to *reduce* pyramidal cell activity in some settings.^44, 88^ We previously demonstrated the pro-excitatory effect, showing activation and altered resonance properties in our isolated PT5B cell model ^98^. However, that model did not show for the observed anti-excitatory effect.

In the current paper, we used a different *I*_h_ model^88^ in our network PT cells, which was able to reconcile these observations: *I*_h_ downregulation reduced PT response to weak inputs, while increasing the cell response to strong inputs ^88, 44, 123, 73^. Furthermore, the pro-excitatory effect seen with strong input, although an isolated change, created widespread activity alterations in the network. We identified stronger activity in the beta/gamma band with reduced *I*_h_ in the resting network (Fig. 6), and increased circuit response to long-range inputs (Figs. 8 and 9).

### 2.5 Preparatory activity vs motor commands

A key question in motor system research is how motor cortex activity gets dissociated from muscle movement during motor planning or mental imagery, and is then shifted to produce commands for action ^38, 120^. One hypothesis has been that this planning-to-execution switch might be triggered by neuromodulation by norepinephrine ^123^. An additional hypothesis is that differential outputs would result from distinct activation of different cells in L5 ^143, 3, 55^. This hypothesis was given further support when a recent study transcriptomically identified different PT subtypes in upper vs lower L5B ^39^. That study also showed that PT5B_upper_ projected to thalamus and generated early preparatory activity, while PT5B_lower_ projected to medulla and generated motor commands.

These two hypotheses are not incompatible, and indeed our simulations demonstrated how these mechanisms might coexist: **1.** Low-ih produced a dramatic increase in PT5B activity, largely triggered by increased excitability of PT with activation via the unidirectional IT to PT projection ^98^. This low-*I*_h_ PT state, associated with norepinephrine activation, would be one factor producing motor action. **2.** Already in the resting condition, the simulation showed markedly different activity in PT5B_upper_ vs PT5B_lower_, with antiphase activity at delta frequency (Fig. 4*D, E*). PT5B_upper_ fired strongly in response to sensory-area inputs, with little PT5B_lower_ activity (Fig. 8). Motor-related inputs more strongly activated activated PT5B_upper_ and produced some PT5B_lower_ activity (Fig. 9). We therefore predict that the planning-to-execution transition might require *both* circuit-level routing of inputs and the neuromodulatory prepared state. The effects of activation via both sensory and motor pathways remains to be tested.

### 2.6 Implications for experimental research and therapeutics

Our model integrates previously isolated experimental data at multiple scales into a unified simulation that can be progressively extended as new data becomes available. This provides a useful tool for researchers in the field, who can use this framework to evaluate hypotheses and guide the design of new experiments using our freely-available model (see Methods). This *in silico* testbed can be systematically probed to study microcircuit dynamics, information flow and biophysical mechanisms with a level of resolution and precision not available experimentally. Unraveling the non-intuitive multiscale interactions occurring in M1 circuits will help us understand disease and develop new pharmacological and neurostimulation treatments for motor disorders ^101, 100, 97, 36, 7, 138, 50, 12, 119^, and improve decoding methods for brain-machine interfaces ^22, 124, 35, 67^.

## 3 Methods

The methods below describe model development with data provenance, and major aspects of the final model. The full documentation of the final model is the source code itself, available for download at http://modeldb.yale.edu/260015.

### 3.1 Morphology and physiology of neuron classes

Seven excitatory pyramidal cell and two interneuron cell models were employed in the network. Their morphology and physiological responses are summarized in Figs. 2*A,B* and 10. In previous work we developed layer 5B PT corticospinal cell and L5 IT corticostriatal cell models that reproduced *in vitro* electrophysiological responses to somatic current injections, including sub- and super-threshold voltage trajectories and f-I curves ^99, 131^. To achieve this, we optimized the parameters of the Hodgkin-Huxley neuron model ionic channels – Na, Kdr, Ka, Kd, HCN, CaL, CaN, KCa – within a range of values constrained by the literature. The corticospinal and corticostriatal cell model morphologies had 706 and 325 compartments, respectively, digitally reconstructed from 3D microscopy images. Morphologies are available via NeuroMorpho.org ^9^ (archive name “Suter Shepherd”). For the current simulations, we further improved the PT model by **1.** increasing the concentration of Ca2+ channels (“hot zones”) between the nexus and apical tuft, following parameters published in ^52^; **2.** lowering dendritic Na+ channel density in order to increase the threshold required to elicit dendritic spikes, which then required adapting the axon sodium conductance and axial resistance to maintain a similar f-I curve; **3.** replacing the HCN channel model and distribution with a more recent implementation ^88^.The new HCN channel reproduced a wider range of experimental observations than our previous implementation ^68^, including the change from excitatory to inhibitory effect in response to synaptic inputs of increasing strength ^44^. This was achieved by including a shunting current proportional to *I*_h_. We tuned the HCN parameters (*lk* and 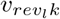) and passive parameters to reproduce the findings noted above, while keeping a consistent f-I curve consistent ^131^.

The network model includes five other excitatory cell classes: layer 2/3, layer 4, layer 5B and layer 6 IT neurons and layer 6 CT neurons. Since our focus was on the role of L5 neurons, other cell classes were implemented using simpler models as a trade-off to enable running a larger number of exploratory network simulations. Previously we had optimized 6-compartment neuron models to reproduce somatic current clamp recordings from two IT cells in layers 5A and 5B. The layer 5A cell had a lower f-I slope (77 Hz/nA) and higher rheobase (250 nA) than that in layer 5B (98 Hz/nA and 100 nA). Based on our own and published data, we found two broad IT categories based on projection and intrinsic properties: corticocortical IT cells found in upper layers 2/3 and 4 which exhibited a lower f-I slope (~72 Hz/nA) and higher rheobase (~281 pA) than IT corticostriatal cells in deeper layers 5A, 5B and 6 (~96 Hz/nA and ~106 pA) ^141, 131, 105^. CT neurons’ f-I rheobase and slope (69 Hz/nA and 298 pA) was closer to that of corticocortical neurons ^105^. We therefore employed the layer 5A IT model for layers 2/3 and 4 IT neurons and layer 6 CT neurons, and the layer 5B IT model for layers 5A, 5B and 6 IT neurons. We further adapted cell models by modifying their apical dendrite length to match the average cortical depth of the layer, thus introducing small variations in the firing responses of neurons across layers.

We implemented models for two major classes of GABAergic interneurons ^51^: parvalbumin-expressing fast-spiking (PV) and somatostatin-expressing low-threshold spiking neurons (SOM). We employed existing simplified 3-compartment (soma, axon, dendrite) models ^69^ and increased their dendritic length to better match the average f-I slope and rheobase experimental values of cortical basket (PV) and Martinotti (SOM) cells (Neuroelectro online database ^134^).

### 3.2 Microcircuit composition: neuron locations, densities and ratios

We modeled a cylindric volume of the mouse M1 cortical microcircuit with a 300 *μm* diameter and 1350 *μm* height (cortical depth) at full neuronal density for a total of 10,073 neurons (Fig. 2). Cylinder diameter was chosen to approximately match the horizontal dendritic span of a corticospinal neuron located at the center, consistent with the approach used in the Human Brain Project model of the rat S1 microcircuit ^85^. Mouse cortical depth and boundaries for layers 2/3, 4, 5A, 5B and 6 were based on our published experimental data ^139, 3, 141^. Although traditionally M1 has been considered an agranular area lacking layer 4, we recently identified M1 pyramidal neurons with the expected prototypical physiological, morphological and wiring properties of layer 4 neurons ^141^ (see also ^14, 10^), and therefore incorporated this layer in the model.

Cell classes present in each layer were determined based on mouse M1 studies ^51, 131, 3, 141, 105, 69, 93^. IT cell populations were present in all layers, whereas the PT cell population was confined to layer 5B, and the CT cell population only occupied layer 6. SOM and PV interneuron populations were distributed in each layer. Neuronal densities (neurons per *mm*^3^) for each layer (Fig. 2*B*) were taken from a histological and imaging study of mouse agranaular cortex ^135^. The proportion of excitatory to inhibitory neurons per layer was obtained from mouse S1 data ^75^. The proportion of IT to PT and IT to CT cells in layers 5B and 6, respectively, were both estimated as 1:1 ^51, 131, 142^. The ratio of PV to SOM neurons per layer was estimated as 2:1 based on mouse M1 and S1 studies ^64, 137^ (Fig. 10*B*). Since data for M1 layer 4 was not available, interneuron populations labeled PV5A and SOM5A occupy both layers 4 and 5A. The number of cells for each population was calculated based on the modeled cylinder dimensions, layer boundaries and neuronal proportions and densities per layer.

**Figure 10:**
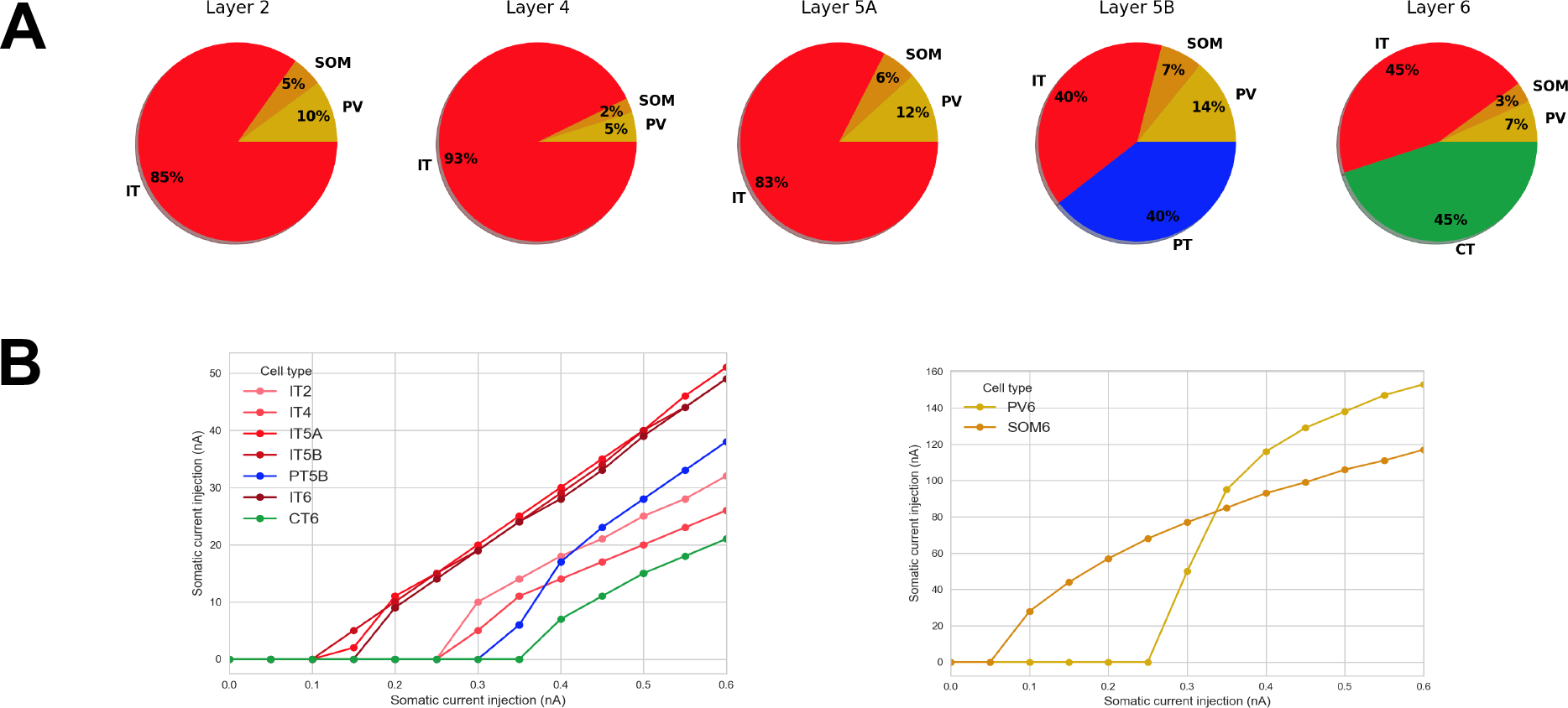
Microcircuit layer composition and cell type f-I response. **A.** Proportion of cell classes per layer; **B.** f-I curve for each excitatory and inhibitory cell types. All properties were derived from published experimental data. Populations labels include the cell class and layer, e.g. ‘IT2’ represents the IT class neurons in layer 2/3.

### 3.3 Local connectivity

We calculated local connectivity between M1 neurons (Figures 2*C* and 11*A*) by combining data from multiple studies. Data on excitatory inputs to excitatory neurons (IT, PT and CT) was primarily derived from mapping studies using whole-cell recording, glutamate uncaging-based laser-scanning photostimulation (LSPS) and subcellular channelrhodopsin-2-assisted circuit mapping (sCRACM) analysis ^139, 3, 141, 142^. Connectivity data was postsynaptic cell class-specific and employed normalized cortical depth (NCD) instead of layers as the primary reference system. Unlike layer definitions which can be interpreted differently between studies, NCD provides a well-defined, consistent and continuous reference system, depending only on two readily-identifiable landmarks: pia (NCD=0) and white matter (NCD=1). Incorporating NCD-based connectivity into our model allowed us to capture wiring patterns down to a 100 *μm* spatial resolution, well beyond traditional layer-based cortical models. M1 connectivity varied systematically within layers. For example, the strength of inputs from layer 2/3 to L5B corticospinal cells depends significantly on cell soma depth, with upper neurons receiving much stronger input ^3^.

**Figure 11:**
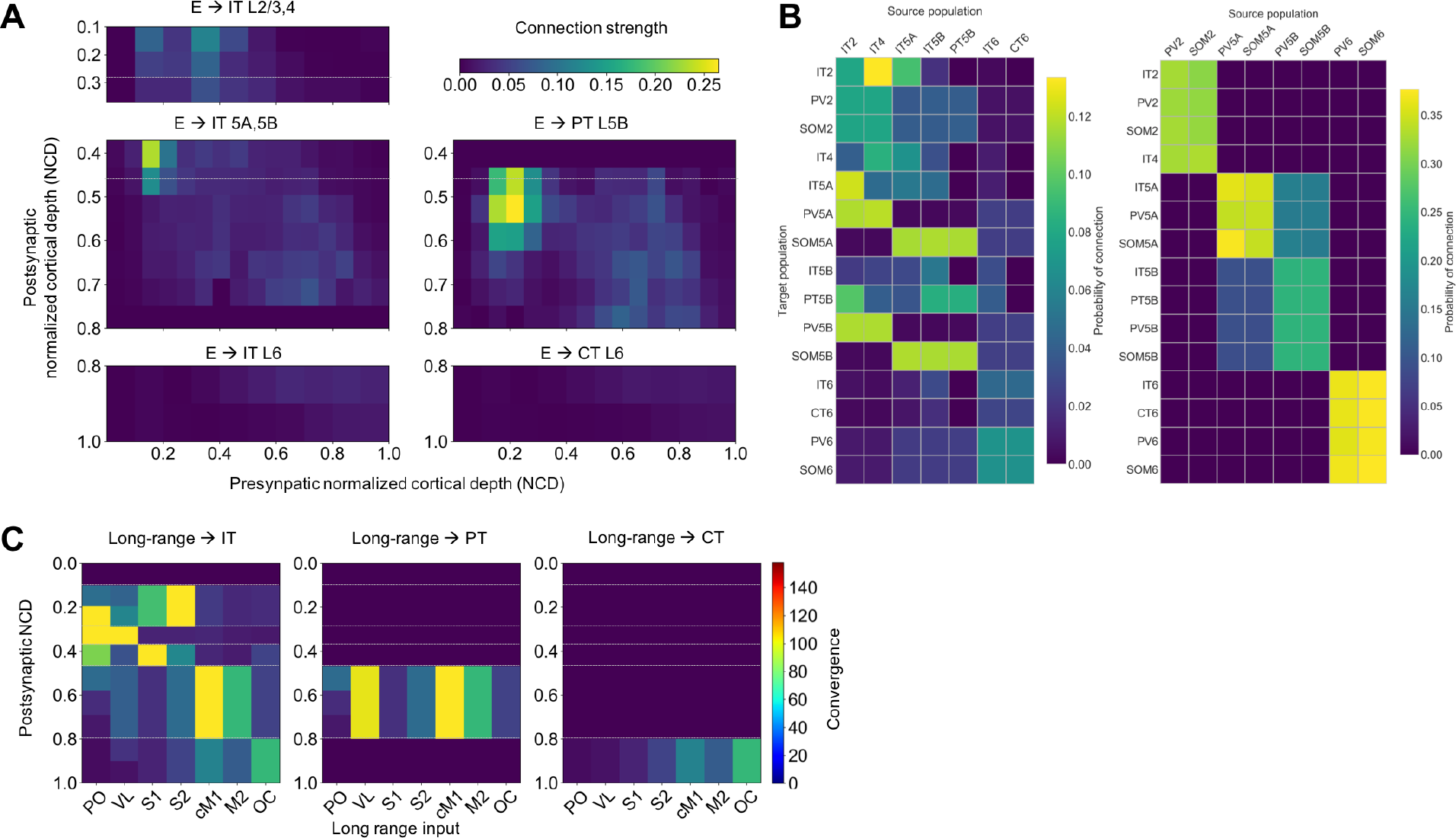
M1 excitatory connectivity: local microcircuitry and and long-range inputs. **A.** Strength of local excitatory connections as a function of pre- and post-synaptic normalized cortical depth (NCD) and post-synaptic cell class; values used to construct the network. **B.** Convergence of long-range excitatory inputs from seven thalamic and cortical regions as a function post-synaptic NCD and cell class; values used to construct the network. **C.** Probability of connection matrix for excitatory (left) and inhibitory (right) populations calculated from an instantiation of the base model network.

Connection strength thus depended on presynaptic NCD and postsynaptic NCD and cell class. For postsynaptic IT neurons with NCD ranging from 0.1 to 0.37 (layers 2/3 and 4) and 0.8 to 1.0 (layer 6) we determined connection strengths based on data from ^139^ with cortical depth resolution of 140 *μm*-resolution. For postsynaptic IT and PT neurons with NCD between 0.37 and 0.8 (layers 5A and 5B) we employed connectivity strength data from ^3^ with cortical depth resolution of 100 *μm*. For postsynaptic CT neurons in layer 6 we used the same connection strengths as for layer 6 IT cells ^139^, but reduced to 62% of original values, following published data on the circuitry of M1 CT neurons ^142^. Our data ^142^ also suggested that connection strength from layer 4 to layer 2/3 IT cells was similar to that measured in S1, so for these projections we employed values from Lefort’s S1 connectivity strength matrix ^75^. Experimentally, these connections were found to be four times stronger than in the opposite direction – from layer 2/3 to layer 4 – so we decreased the latter in the model to match this ratio.

Following previous publications ^66, 75^, we defined connection strength (*s*_*con*_, in mV) between two populations, as the product of their probability of connection (*p*_*con*_) and the unitary connection somatic EPSP amplitude in mV (*v*_*c*_*on*), i.e. *s*_*con*_ = *p*_*con*_ × *v*_*con*_. We employed this equivalence to disentangle the connection *s*_*con*_ values provided by the above LSPS studies into *p*_*con*_ and *v*_*con*_ values that we could use to implement the model. First, we rescaled the LSPS raw current values in pA ^3, 139, 141, 142^ to match *s_con_* data from a paired recording study of mouse M1 L5 excitatory circuits ^66^. Next, we calculated the M1 NCD-based *v*_*con*_ matrix by interpolating a layerwise unitary connection EPSP amplitude matrix of mouse S1 ^75^, and thresholding values between 0.3 and 1.0 mV. Finally, we calculated the probability of connection matrix as *p*_*con*_ = *s*_*con*_/*v*_*con*_.

To implement *v_con_* values in the model we calculated the required NEURON connection weight of an excitatory synaptic input to generate a somatic EPSP of 0.5 mV at each neuron segment. This allowed us to calculate a scaling factor for each segment that converted *v_con_* values into NEURON weights, such that the somatic EPSP response to a unitary connection input was independent of synaptic location. This is consistent with experimental evidence showing synaptic conductances increased with distance from soma, to normalize somatic EPSP amplitude of inputs within 300 *μm* of soma ^82^. Following this study, scaling factor values above 4.0 – such as those calculated for PT cell apical tufts – were thresholded to avoid overexcitability in the network context where each cell receives hundreds of inputs that interact nonlinearly ^128, 11^. For morphologically detailed cells (layer 5A IT and layer 5B PT), the number of synaptic contacts per unitary connection (or simply, synapses per connection) was set to five, an estimated average consistent with the limited mouse M1 data ^56^ and rat S1 studies ^16, 85^. Individual synaptic weights were calculated by dividing the unitary connection weight (*v_con_*) by the number of synapses per connection. Although the method does not account for nonlinear summation effects ^128^, it provides a reasonable approximation and enables employing a more realistic number and spatial distribution of synapses, which may be key for dendritic computations ^78^. For the remaining cell models, all with six compartments or less, a single synapse per connection was used.

For excitatory inputs to inhibitory cell types (PV and SOM) we started with the same values as for IT cell types but adapted these based on the specific connectivity patterns reported for mouse M1 interneurons ^5, 142^ (Fig. 11*A*). Following the layer-based description in these studies, we employed three major subdivisions: layer 2/3 (NCD 0.12 to 0.31), layers 4, 5A and 5B (NCD 0.31 to 0.77) and layer 6 (NCD 0.77 to 1.0). We increased the probability of layer 2/3 excitatory connections to layers 4, 5A and 5B SOM cells by 50% and decreased that to PV cells by 50% ^5^. We implemented the opposite pattern for excitatory connections arising from layer 4,5A,5B IT cells such that PV interneurons received stronger intralaminar inputs than SOM cells ^5^. The model also accounts for layer 6 CT neurons generating relatively more inhibition than IT neurons ^142^. Inhibitory connections from interneurons (PV and SOM) to other cell types were limited to neurons in the same layer ^64^, with layers 4, 5A and 5B combined into a single layer ^93^. Probability of connection decayed exponentially with the distance between the pre- and post-synaptic cell bodies with length constant of 100 *μm* ^43, 42^. We introduced a correction factor to the distance-dependent connectivity measures to avoid the *border effect*, *i.e.,* cells near the modeled volume edges receiving less or weaker connections than those in the center.

For comparison with other models and experiments, we calculated the probability of connection matrices arranged by population (instead of NCD) for the base model network instantiation used throughout the results. (11*B*).

Excitatory synapses consisted of colocalized AMPA (rise, decay *τ*: 0.05, 5.3 ms) and NMDA (rise, decay *τ*: 15, 150 ms) receptors, both with reversal potential of 0 mV. The ratio of NMDA to AMPA receptors was 1.0 ^92^, meaning their weights were each set to 50% of the connection weight. NMDA conductance was scaled by 1/(1 + 0.28 · *Mg* · exp (−0.062 *· V*)); Mg = 1mM ^61^. Inhibitory synapses from SOM to excitatory neurons consisted of a slow *GABA_A_* receptor (rise, decay *τ*: 2, 100 ms) and *GABA*_*B*_ receptor, in a 90% to 10% proportion; synapses from SOM to inhibitory neurons only included the slow *GABA_A_* receptor; and synapses from PV to other neurons consisted of a fast *GABA_A_* receptor (rise, decay *τ*: 0.07, 18.2). The reversal potential was −80 mV for *GABA_A_* and −95 mV for *GABA_B_*. The *GABA_B_* synapse was modeled using second messenger connectivity to a G protein-coupled inwardly-rectifying potassium channel (GIRK) ^33^. The remaining synapses were modeled with a double-exponential mechanism.

Connection delays were estimated as 2 ms plus a variable delay depending on the distance between the pre- and postsynaptic cell bodies assuming a propagation speed of 0.5 m/s.

### 3.4 Long-range input connectivity

We added long-range input connections from seven regions that are known to project to M1: thalamic posterior nucleus (PO), ventro-lateral thalamus (VL), primary somatosensory cortex (S1), secondary somatosensory cortex (S2), contralateral primary motor cortex (cM1), secondary motor cortex (M2) and orbital cortex (OC). Each region consisted of a population of 1000 ^28, 16^ spike-generators (NEURON VecStims) that generated independent random Poisson spike trains with uniform distributed rates between 0 and 5 Hz ^140, 54^; or 0 to 10 Hz ^58, 60^ when simulating increased input from a region. Previous studies provided a measure of normalized input strength from these regions as a function of postsynaptic cell type and layer or NCD. Broadly, PO ^141, 142, 93^, S1 ^83^ and S2 ^132^ projected strongly to IT cells in layers 2/3 and 5A (PO also to layer 4); VL projected strongly to PT cells and to layer 4 IT cells ^141, 142, 93^; cM1 and M2 projected strongly to IT and PT cells in layers 5B and 6 ^55^; and OC projected strongly to layer 6 CT and IT cells ^55^. We implemented these relations by estimating the maximum number of synaptic inputs from each region and multiplying that value by the normalized input strength for each postsynaptic cell type and NCD range. This resulted in a convergence value – average number of synaptic inputs to each postsynaptic cell – for each projection Fig. 11*C*. We fixed all connection weights (unitary connection somatic EPSP amplitude) to 0.5 mV, consistent with rat and mouse S1 data ^56, 28^.

To estimate the maximum number of synaptic inputs per region, we made a number of assumptions based on the limited data available (Fig. 11*C* and 2*B*). First, we estimated the average number of synaptic contacts per cell as 8234 by rescaling rat S1 data ^87^ based on our own observations for PT cells ^131^ and contrasting with related studies ^122, 31^; we assumed the same value for all cell types so we could use convergence to approximate long-range input strength. We assumed 80 % of synaptic inputs were excitatory vs. 20 % inhibitory ^31, 85^; out of the excitatory inputs, 80 % were long-range vs. 20 % local ^85, 129^; and out of the inhibitory inputs, 30 % were long-range vs. 70 % local ^129^. Finally, we estimated the percentage of long-range synaptic inputs arriving from each region based on mouse brain mesoscale connectivity data ^104^ and other studies ^86, 16, 87, 146, 14^.

### 3.5 Dendritic distribution of synaptic inputs

Experimental evidence demonstrates the location of synapses along dendritic trees follows very specific patterns of organization that depend on the brain region, cell type and cortical depth ^108, 132^; these are likely to result in important functional effects ^72, 74, 128^. We employed sCRACM data to estimate the synaptic density along the dendritic arbor – 1D radial axis – for inputs from PO, VL, M2 and OC to layers 2/3, 5A, 5B and 6 IT and CT cell ^55^ (Fig. 12*A*), and from layer 2/3 IT, VL, S1, S2, cM1 and M2 to PT neurons ^132^ (Fig. 12B). To approximate radial synaptic density we divided the sCRACM map amplitudes by the dendritic length at each grid location, and averaged across rows. Once all network connections had been generated, synaptic locations were automatically calculated for each cell based on its morphology and the pre- and postsynaptic cell type-specific radial synaptic density function (Fig. 12C). Synaptic inputs from PV to excitatory cells were located perisomatically (50 *μm* around soma); SOM inputs targeted apical dendrites of excitatory neurons ^93, 64^; and all inputs to PV and SOM cells targeted apical dendrites. For projections where no data synaptic distribution data was available – IT/CT, S1, S2 and cM1 to IT/CT cells – we assumed a uniform dendritic length distribution.

**Figure 12:**
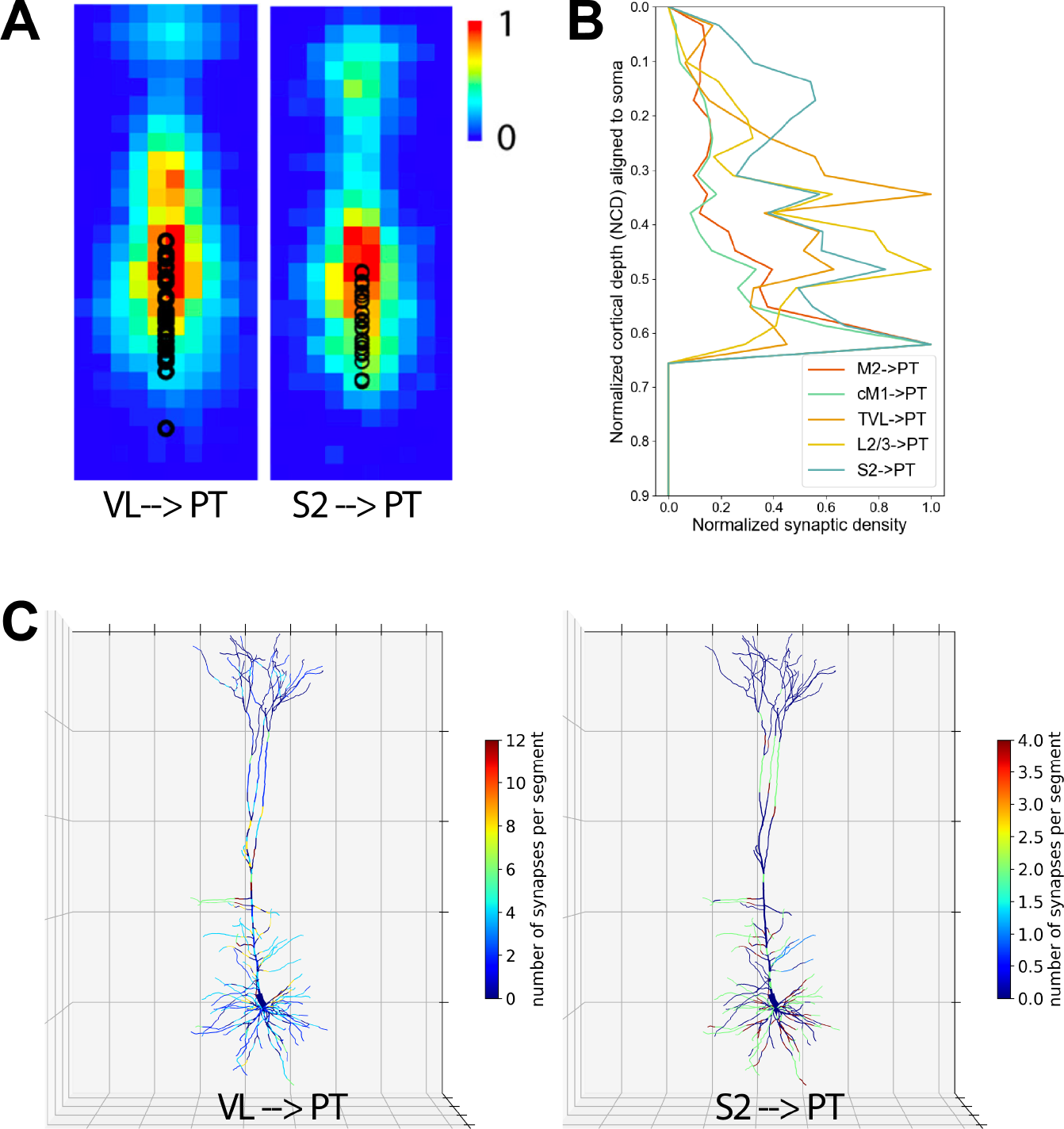
Dendritic distribution of synaptic inputs. **A.** Average 2D sCRACM maps for VL →PT and S2 →PT inputs (experimental data). **B.** Synaptic density profile (1D) along the dendritic arbor for inputs from layer 2/3 IT, VL, S1, S2, cM1 and M2 to PT neurons. Calculated by normalizing sCRACM maps by dendritic length at each grid location and averaging across rows. **C.** Synaptic density per neuron segment automatically calculated for each neuron based on its morphology and the pre- and postsynaptic cell type-specific radial synaptic density function. Here, VL →PT and S2 →PT are compared and exhibit partially complementary distributions.

### 3.6 Model implementation, simulation and analysis

The model was developed using parallel NEURON (neuron.yale.edu)^80^ and NetPyNE (www.netpyne.org)^37^, a Python package to facilitate the development of biological neuronal networks in the NEURON simulator. NetPyNE emphasizes the incorporation of multiscale anatomical and physiological data at varying levels of detail. It converts a set of simple, standardized high-level specifications in a declarative format into a NEURON model. This high-level language enables, for example, defining connectivity as function of NCD, and distributing synapses across neurons based on normalized synaptic density maps. NetPyNE facilitates running parallel simulations by taking care of distributing the workload and gathering data across computing nodes, and automates the submission of batches of simulations for parameter optimization and exploration. It also provides a powerful set of analysis methods so the user can plot spike raster plots, LFP power spectra, information transfer measures, connectivity matrices, or intrinsic time-varying variables (eg. voltage) of any subset of cells. To facilitate data sharing, the package saves and loads the specifications, network, and simulation results using common file formats (Pickle, Matlab, JSON or HDF5), and can convert to and from NeuroML ^47, 46^ and SONATA ^30^, standard data formats for exchanging models in computational neuroscience. Simulations were run on XSEDE supercomputers Comet and Stampede, using the Neuroscience Gateway (NSG) and our own resource allocation, and on Google Cloud supercomputers.

NetPyNE facilitates optimization and exploration of network parameters through automated batch simulations. The user specifies the range of parameters and parameter values to explore and the tool automatically submits the jobs in multicore machines (using NEURON’s Bulletin board) or HPCs (using SLURM/Torque). Multiple pre-defined batch simulation setups can be fully customized for different environments. We ran batch simulations using NetPyNE’e automated SLURM job submission on San Diego Supercomputer Center’s (SDSC) Comet supercomputer and on Google Cloud Platform.

The NetPyNE tool also includes the ability to simulate local field potentials (LFPs) obtained from extracellular electrodes located at arbitrary 3D locations within the network. The LFP signal at each electrode is obtained using the “line source approximation” ^107, 19, 77^, which is based on the sum of the membrane current source generated at each cell segment divided by the distance between the segment and the electrode. The calculation assumes that the electric conductivity and permittivity of the extracellular medium are constant everywhere and do not depend on frequency.

Spectral Granger Causality analysis (Fig. 7) was also performed via the NetPyNE package, which follows the formulation described in ^62^ using a Python implementation adapted from the Matlab BSMART package ^29^. This implementation of Spectral Granger Causality has previously bee used to analyze spiking network models ^65^.

Modulation index (Fig. 4) was used as a measure of cross-frequency coupling between the phase of a slow frequency and the amplitude (or envelope) of a faster frequency. The modulation index calculation was implemented in Python based on the method described in ^133^ and making use of a set of functions to filter the fast frequency at variable bandwidths ^13^.

To study the significance of spectral Granger casuality and modulation index (Figs. 4*E* and 7*F*) results we generated 50 shuffled versions of the input data and then calculated the corresponding surrogate measures (spectral Granger causality or modulation index), from which we could infer the chance distribution. We then calculated the z-score of the original simulation results, which indicates the number of standard deviations from the shuffled sample mean. Z-scores can be directly related to p-values, e.g., assuming a one-tailed hypothesis a z-score of 3.29 corresponds to a p-value *≤* 0.0005.

## 4 Acknowledgements

This work was funded by the following grants: NIH U01EB017695 (WWL), NYS SCIRB DOH01-C32250GG-3450000 (SDB,WWL), NIH R01EB022903 (WWL), NIH U24EB028998 (SDB), NIH R01DC012947 (SAN), ARO W911NF-19-1-0402 (SAN). The views and conclusions contained in this document are those of the authors and should not be interpreted as representing the official policies, either expressed or implied, of the U.S. Government or any of its agencies. The U.S. Government is authorized to reproduce and distribute reprints for Government purposes notwithstanding any copyright notation herein.

